# The Effect of Sunlight Intensity on Flower Opening Time and Exposure Duration in Rice (*Oryza sativa* ssp. *indica*) Landraces

**DOI:** 10.1101/2023.09.10.557080

**Authors:** Debal Deb, Sreejata Dutta, Mahendra Nauri

## Abstract

Records of sunlight intensity at anthesis of 388 rice landraces (with 32 replicated populations), flowering in the short-day season of 2022, reveal that under cloudy condition, the rice florets tend to open later, or the sunrise-to-anthesis duration (SAD) is longer when rice florets open than at sunny period. This difference in the length of SAD was statistically highly significant (*p* < 0.0001), confirmed by two sample permutation test with 10,000 iterations. This finding corroborates our general observation previously reported from a larger set of 1114 landraces (including the 388 landraces in this study). However, the intensity of sunlight at the flower opening time (FOT) may not remain uniformly sunny (high illuminance) or cloudy (low illuminance, < 40000 lux) until the flower closing time (FCT). To understand the effect of uniformly low sunlight intensity, we subsequently recorded solar illuminance at FOT of 33 landrace populations (including 8 repeats). Half of each population was kept under artificial shade, compared to the other half exposed to sunlight. This experiment revealed that low illuminance, mimicking overcast days, significantly (*p* < 0.01) delays FOT and lengthens SAD, corroborating the pattern detected in our earlier findings. Permutation tests with 10,000 iterations decisively confirms (*p* < 0.0001) the prolongation of SAD under shade and cloudy condition. Experimental shading has an indeterminate effect on FED of the same landraces. We surmise that the delayed FOT during natural cloudy period is an adaptation in rice plants in anticipation of rain, for protection of the pollen from rainwash.

## 1. Introduction

Reproductive biology of rice (*Oryza sativa*) is primarily based on the anthesis date and time, and different rice landraces are known to open usually 1 day after booting. The rice flower opens to facilitate stamen exsertion and pollen release ^[1,2]^, leading to the pollination of the flowers of neighbouring rice plants. In spite of the received wisdom ^[1–3]^ that cross pollination frequency in rice rarely exceeds 2%, cross pollination is likely to occur at a high frequency between neighbouring cultivars with close proximity of flower opening time (FOT) and a large temporal overlap of the duration of flowers remaining open (=flower exposure duration, FED) ^[4]^. Despite the crucial importance of FOT and FED in cross pollination required in rice breeding, there is hardly any record of FED overlaps between the crossing parents in published literature on experimental cross pollination in rice, until recently ^[5]^. The FOT and FED are known to be different for different cultivars, and influenced by various environmental factors ^[6–8]^.

Our records of FOT and FED in 1114 *indica* landraces, constituting the largest database of rice anthesis ^[9]^ suggests that a majority of these landraces is likely to be cross pollinated at a high rate due to considerable overlaps of FOT and FED among hundreds of landraces. However, the FOT and FED of the same cultivar may vary with seasonal photoperiod (short day vs long day), daylight condition (sunny vs cloudy)^[2]^, and day temperature^[6-8, 10]^. The first study of the variability of FOT and FED with flowering season and daylight conditions^[5]^ indicated that an adequate solar illuminance is likely as important to elicit FOT as day maximum temperature. We observed that the anthesis time of most landraces recorded on sunny days considerably changed on cloudy days. However, the ‘cloudy’ hours included the periods of cloud cover with intermittent sunshine, as well as fully overcast periods at FOT, and therefore a more precise description of ‘cloudiness’ at the time of anthesis is required to examine the influence of sunlight intensity on rice flower opening and closing behaviour. This led us to explore the effect of incident sunlight intensity at the time of anthesis on both FOT and FED. The present study describes our attempt to find a relationship between sunlight intensity with FOT and FED, based on empirical observations of flowering times and durations of 390 rice landraces (420 populations), and on a comparison of 25 landraces subjected to full exposure to daylight and under shading (mimicking cloudy days) in a field experimental setup during the same season.

Detecting any difference between FEDs of landraces exposed to normal solar illuminance and those under cloudy/ shaded condition would provide an important guideline for rice breeding experiments, in order to choose the appropriate landraces with adequate overlap of FED. Our findings would also be useful in maintaining genetic purity of selected rice landraces, by ensuring zero overlap of FED between the neighbouring cultivar populations.

## 2. Materials and Methods

### 2.1 Study site, cultivation season and materials

We documented the FOT and FED of 388 landraces grown on Basudha conservation farm (http://cintdis.org/basudha), located in Rayagada district of southern Odisha (19°42′32.0″N, 83°28′8.4″E) during short-day (*Aman*) season of the year 2022. Some of the 388 landraces were replicate, so the tota number landrace populations was 420, each population planted in blocks of 8 rows × 8 columns. The procedure of recording of FOT, flower closing time (FCT), FED (the difference between FOT and FCT), and sunrise to anthesis duration (SAD) on the days of flowering of each of these landraces is given in our previous works^[5, 8]^. In brief, our methods of observation and recording were as follows.

a. Two of us (DD and MN) stood before each landrace population on the day of anthesis from 8:40 am, to record the exact time of opening of an apical floret in one of the first 50% exserted panicles in each landrace population (consisting of 64 plants), and tagged the stalk of the panicle with a coloured thread, without touching any florets. Among the selected panicles, we recorded the FOT of the floret that opened the earliest. This procedure was repeated in each varietal plot.
b. We also recorded the time of full closure of the last floret in each population. The FED was calculated as the time interval between the FOT of the first floret and the FCT of the last floret in a varietal plot.
c. We recorded the changes in FOT on sunny and cloudy days. If the florets of a selected panicle opens and/ or closes on a cloudy day, we recorded both the FOT and FCT of other florets on a different panicle of the same cultivar on a subsequent sunny day. This was not possible for every cultivar, so the number of observations of FOT and FCT on sunny and cloudy days were not equal.
d. We estimated the sunrise to anthesis duration (SAD) as the interval between sunrise time and FOT. The local sunrise time on each day of anthesis was obtained mainly from https://www.timeanddate.com/sun/@10775335 and https://isubqo.com/prayer-time/india/odisha/bishama-katek.
e. We further recorded the intensity of sunlight incident on the panicles at FOT and FCT of each of the cultivar population during short-day (*Aman*) season of 2022 (**Table 1**). For measuring incident sunlight illuminance, we used a Digital Illuminance Meter (http://drmeter.com/products/lx1330b-digital-illuminance-light-meter, model No. LX1330B) held next to the panicle, without touching it, at the onset of florets opening and after the full closure of the last florets on the panicle. The maximum solar illuminance (Lmax) at FOT incident at the panicle level on all cloudy days never exceeded 40K lux, and therefore we decided on a threshold Lmax of “sunny” hours to be 40 K lux. Although all flowers opened and closed at daytime solar wavebands between 400 nm to 700 nm wavelength, we chose not to convert solar illuminance in lux into photosynthetically active radiation (PAR) in μmol of photons/m^2^/s because photosynthetic activity per se was often limited on cloudy days, especially during cloudy hours, and severely truncated under experimental shade.

**Table 1.** Cultivation Seasons and Phenological Stages of 390 Landraces Examined.

| Season | No. of Landraces (excl. repeats) | Sowing Dates | Flowering Dates (FD) | Dates of Harvest | No. of Populations (incl. repeats) |
| --- | --- | --- | --- | --- | --- |
| *Aman* 2022 | 388 | 21 Jun - 06 Jul | 31 Aug - 27 Nov | 25 Sep - 07 Feb | 420 |

### 2.2 Field experiment

During the same short-day (*Aman*) season of 2022, we set up an experiment with 26 landraces in the same field to compare the FOT and FED of these landraces under natural illuminance in open sunlight and low illuminance under shade. Light intensity was measured during the FOT of the first florets and the FCT of the last florets of each landrace population in this experiment, by lux meters, as described above.

The design of the field experiment is shown in **Fig. 1**. An opaque polythene sheet was built over half of each landrace population, and kept in place from the booting stage until after the last FCT of the population. The shade allowed entry of only diffused sunlight from the sides, with incident light intensity never exceeding 5825 lux (Table S-1), mimicking overcast and rainy days. This portion of the landrace population kept in dark under shade was designated “population SH”, while the other half of the same population was left fully exposed to sunlight (designated “population OL”). The Lmax incident on the population OL during the study period exceeded 30000 lux throughout the study period.

**Fig 1:**
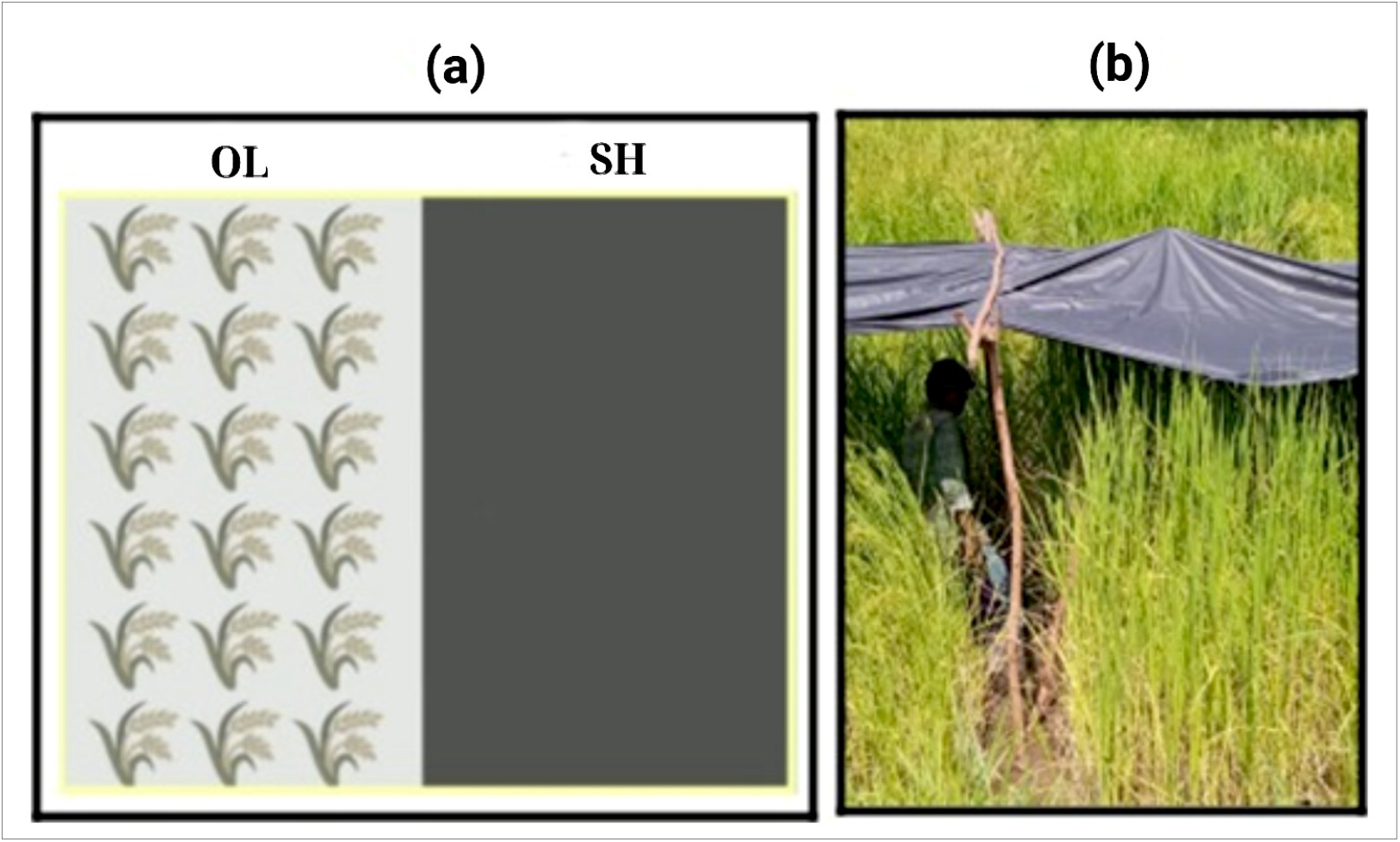
**(a)** The Experiment with Illuminance. Half of each illustrative landrace population was exposed to open sunlight (OL), while the other half was kept under shade (SH) of an opaque black polythene sheet. **(b)** Reading lux meter next to a flowering panicle under the shade.

### 2.3 Statistical analyses

Data sorting, time data conversion, and regression analyses were made using Open Office software on a laptop computer. The data distribution was evaluated using summary statistics and box plots with R statistical software version 4.3.0. For visual assessment of normality, we used QQ plots and performed Shapiro-Wilk and/or Anderson-Darling tests, both online ^[11]^ and with R. For data sets where the QQ-plot showed the data distribution approaching normality, we performed ANOVA, and conducted a thorough residual analysis. When the ANOVA results indicated significant differences in means among groups, we employed Tukey’s Honest Significant Difference (HSD) test to conduct pairwise comparisons between these groups.

For data whose distribution violated the assumption of normality, ANOVA and *t* tests were not performed.^12]^ Instead, a non-parametric approach with Wilcoxon-Mann-Whitney test ^[12]^ was employed to estimate the significance of the difference in SAD and FED medians of the populations between sunny and cloudy/ shaded conditions. The hypotheses considered in this analyses are:

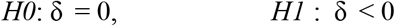

where δ is the difference between medians of SAD-S and SAD-C (or between the medians of FED-S and FED-C). We chose the confidence interval at 95% for all statistical analyses. To control for multiple testing, we used Bonferroni corrections to draw valid conclusions. We also conducted a two-sample permutation test ^[11]^ for location, considering the same hypotheses as above, and conducted 10,000 iterations for the 25 paired data. All analyses were conducted using R statistical software version 4.3.0.

To assess the effect of sunlight intensity, we plotted the SAD and FED on Lmax, and estimated the coefficient of determination (*R*^2^) and *t* to determine if the slope of regression was significantly different from zero.

## 3. Results

We describe the results in two sections, namely, the results of analyses of the relationship of FOT, SAD and SED with different daylight conditions on the day of anthesis of 420 landrace populations in section *3*.*1*; and the same analyses for the selected 33 populations experimentally kept under shade as well as exposed to open sunlight, in section *3*.*2*.

### 3.1. General field observations

The data of FOT and FED, as well as the day temperatures and the daylight conditions on the respective flowering dates of each of 420 landraces grown in 2022 are available on Harvard Dataverse^[9]^. In addition to these data, the sunlight intensity at the respective FOT and FCT of each of the total of 420 landrace populations are deposited with Harvard Dataverse^[14]^. On a few overcast and rainy days, the minimum solar intensity was often less then 3.5 K lux. For instance, on the 14th September, 20th September and the 21st October 2022, minimum illuminance at FOT were 2399 lux, 3343 lux, and 2915 lux, respectively. On intermittently cloudy days, the Lmax never exceeded 40 K lux. By contrast, during sunny periods, MSL always exceeded 40000 lux, reaching up to 182 K lux^[14].^

#### 3.1.1. The effect of daylight conditions on SAD and FED

The landraces tend to open their florets considerably earlier (*n* = 239, mean SAD = 255.9 min, median = 256 min., IQR = 22) on sunny days (light intensity ≥ 40 K lux) than on cloudy days with sunlight intensity <40 K lux (*n* = 181, mean SAD = 283, median = 280 min., IQR = 31) during the same season. The mean values of SAD and FED exposed to full sunlight and under cloudy conditions are shown in **Fig. 2**. The box plot (Supplementary **Fig. S1**) and the QQ plot (Supplementary **Fig. S2**) indicate that the distribution of SAD, under both sunny and cloudy conditions, approaches normality. Although two-sample *t*-test reveals a significant difference (*t* = 11.26, *p* <0001) between the two groups, we decided to ignore this result. This decision was based on the Shapiro-Wilk test, which indicated a slight deviation from normality due to the presence of heavy tails. We therefore performed Wilcoxon-Mann-Whitney test, which showed a highly significant (*z* = 10.73, Bonferroni adjusted *p* < 0.0001) difference between the median values on sunny and cloudy periods. Permutation test of the same data with 10,000 iterations for location decisively confirms that the difference was highly significant (*p* < 0.0001) (**Table 3**, part **A**). Between the FED of the landraces during sunny period (*n =* 239, mean FED = 72 min., median = 68 min., IQR = 40.5) and during cloudy period (*n* = 181, mean FED = 69 min., median = 66 min, IQR =22), no statistical difference was detected (Wilcoxon-Mann-Whitney *z* = – 0.6901, Bonferroni adjusted *p* = 0.98). Two-sample permutation test with 10,000 iterations for location decisively confirmed the same result (*p* = 0.87).

**Fig 2:**
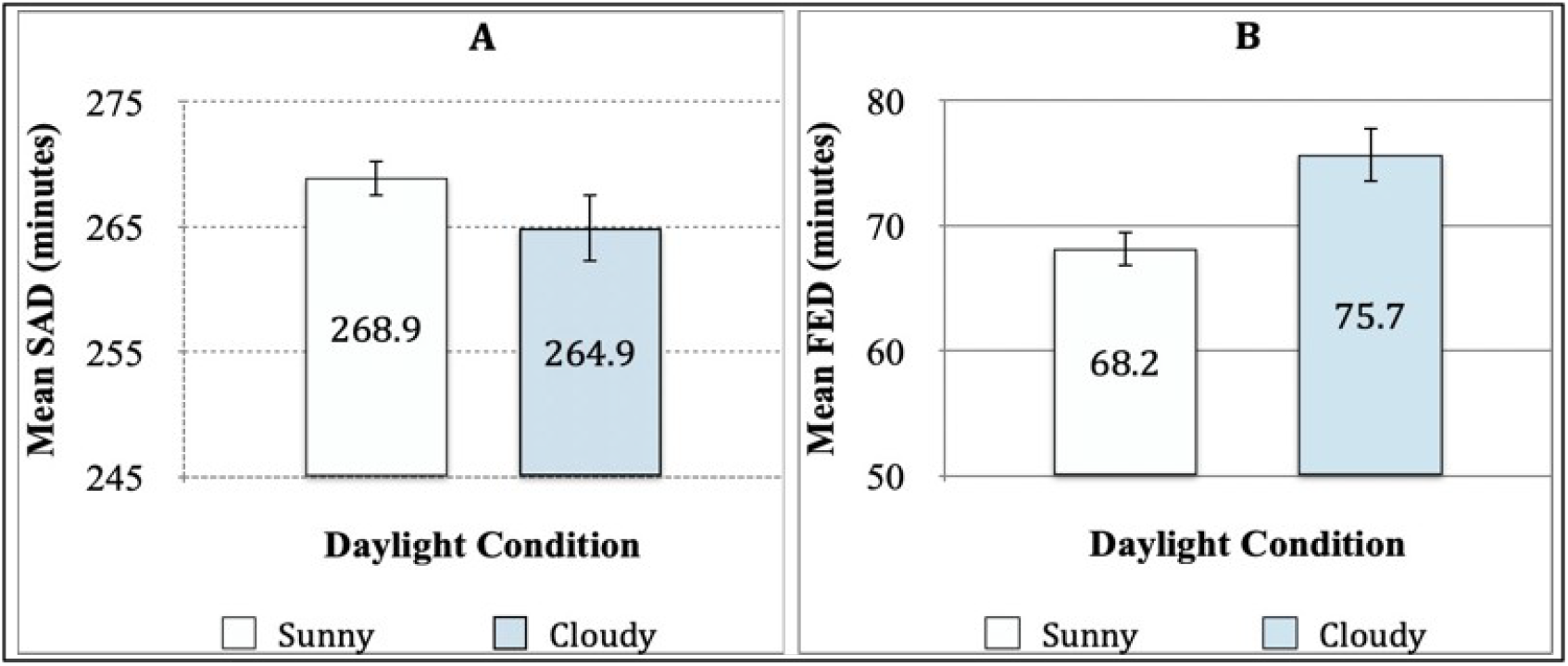
Mean Values**(A)** SAD (minutes) and **(B)** FED (minutes) on Sunny (*n* = 300) and Cloudy Days (*n* = 120). Vertical bars show the standard errors.

The validity of our classification of “sunny” (S) and “cloudy” (C) conditions, based on the threshold value of illuminance (40K lux) in the field condition, is endorsed by an overall power function relationship between SAD and Lmax for all the landrace populations (*n* = 420):

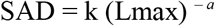

with constant k = 353.98 and exponent *a* = 0.71. This relationship is highly significant (*R*^2^ = 0.23, *t* = 10.49, *p* <0.001), indicating a prominent trend of SAD declining with greater illuminance at FOT (**Fig. 3**).

**Fig 3:**
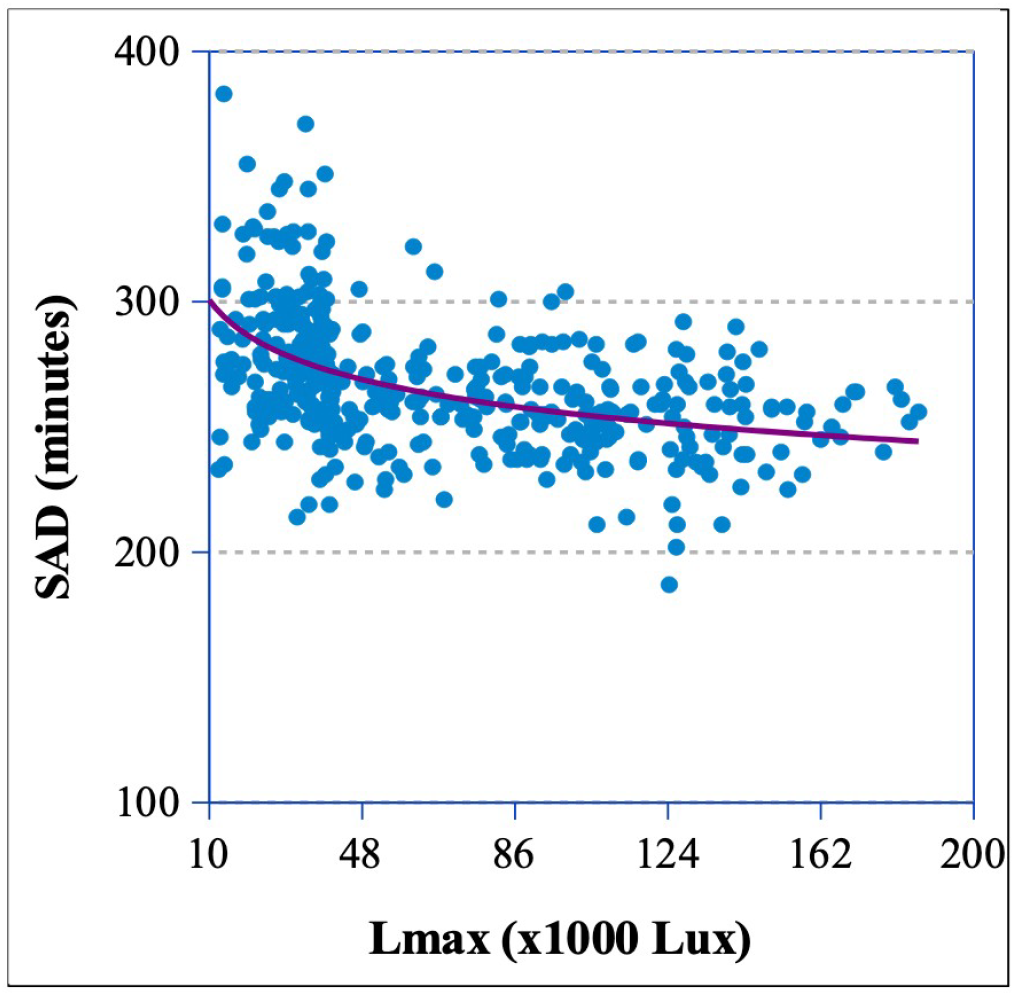
Power function relationship of SAD with Lmax, SAD = 353.98 (Lmax) − ^0.71^

#### 3.1.2. The relationship between FOT/ SAD and FED during sunny and cloudy periods

When the daylight conditions (S or C) exactly at FOT are examined as separate groups, FED shows a strong inverse relationship with FOT (**Fig. 3A**), and equivalently, with SAD (**Fig. 3B**). The coefficient of determination (*R*^2^) for FED on both FOTd and SAD are almost identical, and indicates that delayed anthesis is strongly associated with shorter FED in landraces.

#### 3.1.3. The effect of solar illuminance at FOT, combined with illuminance at FCT

To examine if the inverse relationship between FOT and FED holds for these landraces closing their flowers at different sunlight exposures, we classified the phenological observations into four sunlight exposure categories (**Table 2**), namely, flowers opening and closing at sunny hours (SS, 158 landraces, 161 populations); opening at sunny but closing at cloudy period (SC, 77 landraces, 78 populations); opening and closing both at cloudy periods (CC, 66 landraces, 68 populations); and opening at cloudy but closing when it was sunny (CS, 108 landraces, 113 populations). Because two or more replicated populations of a given landrace may belong in the same or different categories, the count of the landraces from SS and SC categories combined (SS+ SC = 235) exceeded the actual number of landraces opening their flowers at S periods without repeats (= 226). For the same reason, the count of the landraces from CS and CC categories (CS+ CC = 174) exceeds the actual number of landraces opening their flowers at C periods without repeats (= 173), as shown in **Table 2**.

**Table 2.** Schema of recording of incident sunlight intensity at FOT and FCT of 420 landrace populations (including replications). Numbers in parentheses are the number of landraces (excluding repetitions) in each category.

| Sunlight at FOT | No. of Populations<br>(No. of Landraces<br>without repeat) | Sunlight at FCT | Category | No. of Populations<br>(No. of Landraces<br>without repeat) |
| --- | --- | --- | --- | --- |
| Sunny<br>(>40,000 lux) | 239<br>(226) | *At Sunny hour* | SS | 161<br>(158) |
|  |  | *At Cloudy hour* | SC | 78<br>(77) |
| Cloudy<br>(≤40,000 lux) | 181<br>(173) | *At Sunny hour* | CS | 113<br>(108) |
|  |  | *At Cloudy hour* | CC | 68<br>(66) |
| *Total* | 420<br>(388) |  |  |  |

**Table 3.**
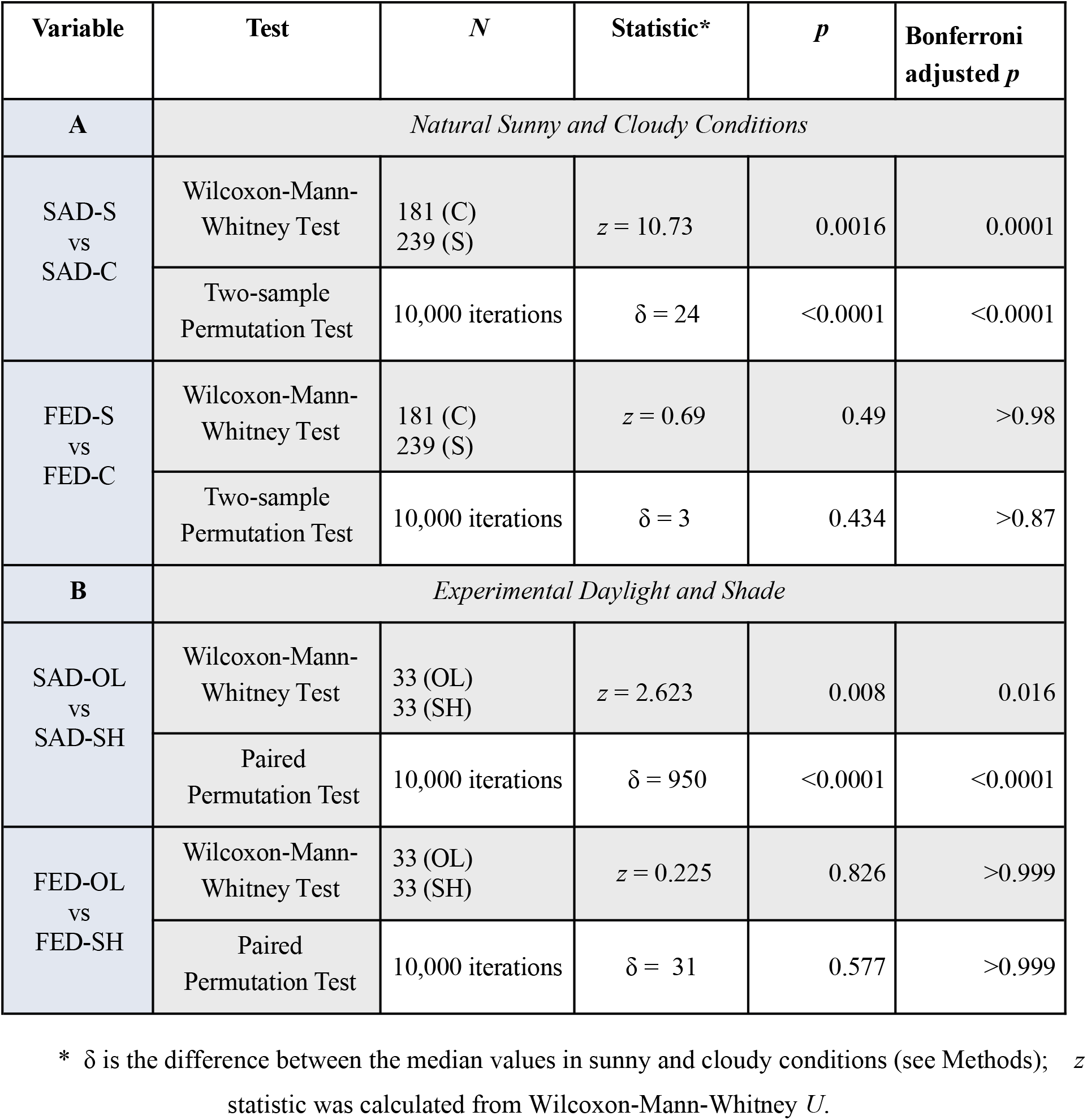
Statistical Tests of Significance [Part **A**] between 181 Landrace Populations in Cloudy (C) and 239 Populations in Sunny (S) Periods at Anthesis, and [Part **B**] between 33 pairs of Landrace Populations Under Open Sun (OL) and under Shade (SH).

| Variable | Test | *N* | Statistic* | *p* | Bonferroni adjusted *p* |
| --- | --- | --- | --- | --- | --- |
| **A** | *Natural Sunny and Cloudy Conditions* |  |  |  |  |
| SAD-S<br>vs<br>SAD-C | Wilcoxon-Mann-Whitney Test | 181 (C)<br>239 (S) | $z = 10.73$ | 0.0016 | 0.0001 |
| | Two-sample Permutation Test | 10,000 iterations | $\delta = 24$ | <0.0001 | <0.0001 |
| FED-S<br>vs<br>FED-C | Wilcoxon-Mann-Whitney Test | 181 (C)<br>239 (S) | $z = 0.69$ | 0.49 | >0.98 |
| | Two-sample Permutation Test | 10,000 iterations | $\delta = 3$ | 0.434 | >0.87 |
| **B** | *Experimental Daylight and Shade* |  |  |  |  |
| SAD-OL<br>vs<br>SAD-SH | Wilcoxon-Mann-Whitney Test | 33 (OL)<br>33 (SH) | $z = 2.623$ | 0.008 | 0.016 |
| | Paired Permutation Test | 10,000 iterations | $\delta = 950$ | <0.0001 | <0.0001 |
| FED-OL<br>vs<br>FED-SH | Wilcoxon-Mann-Whitney Test | 33 (OL)<br>33 (SH) | $z = 0.225$ | 0.826 | >0.999 |
| | Paired Permutation Test | 10,000 iterations | $\delta = 31$ | 0.577 | >0.999 |

\* $\delta$ is the difference between the median values in sunny and cloudy conditions (see Methods); $z$ statistic was calculated from Wilcoxon-Mann-Whitney $U$ .

The relationship of FED with FOTd for the populations under SS, SC, CS and CC conditions is inverse, and all are significantly (*p* <0.01) different from zero (**Fig. 5**). This pattern of relationship is corroborative of the strongly inverse (*p* < 0.0001) relationship of FED with FOTd of the same populations, when they are examined at S and C conditions at FOT alone (**Fig. 4a**).

**Fig 4:**
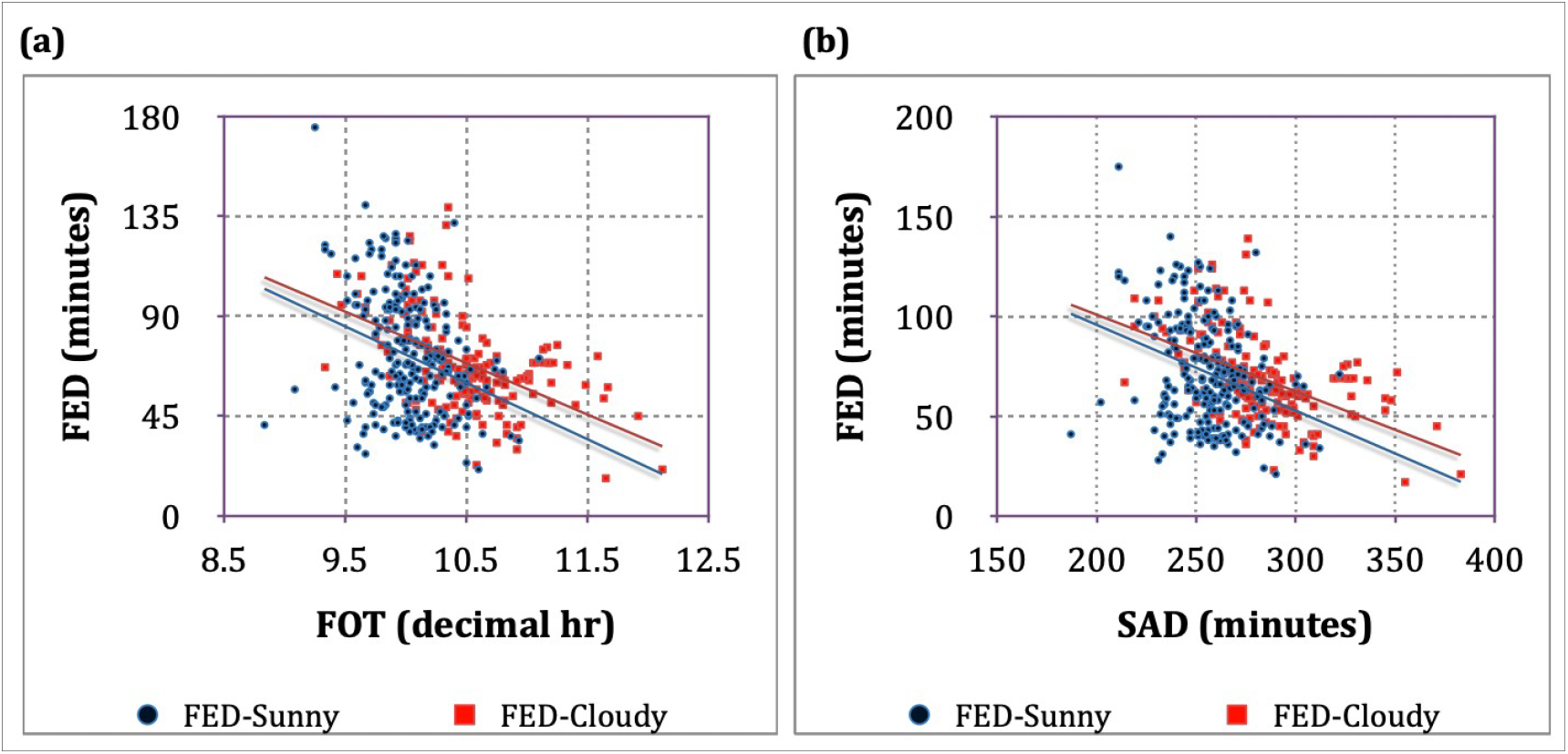
The regression of FED **(a)** on FOT (on Sunny days, *df* = 239, *b* = – 25.42, *R*^2^ = 0.09, *t* = 4.75; on Cloudy days, *df* = 181, *b* = – 23.17, *R*^2^ = 0.26, *t* = 7.4); and **(b)** on SAD (for Sunny days, *df* = 239, Slope *b* = – 0.429, *R*^2^ = 0.09, *t* = 4.72; for Cloudy days, *df* = 181, *b* = – 0.384, *R*^2^ = 0.25, *t* = 7.23); *p* < 0.00001 for all slopes.

**Fig 5:**
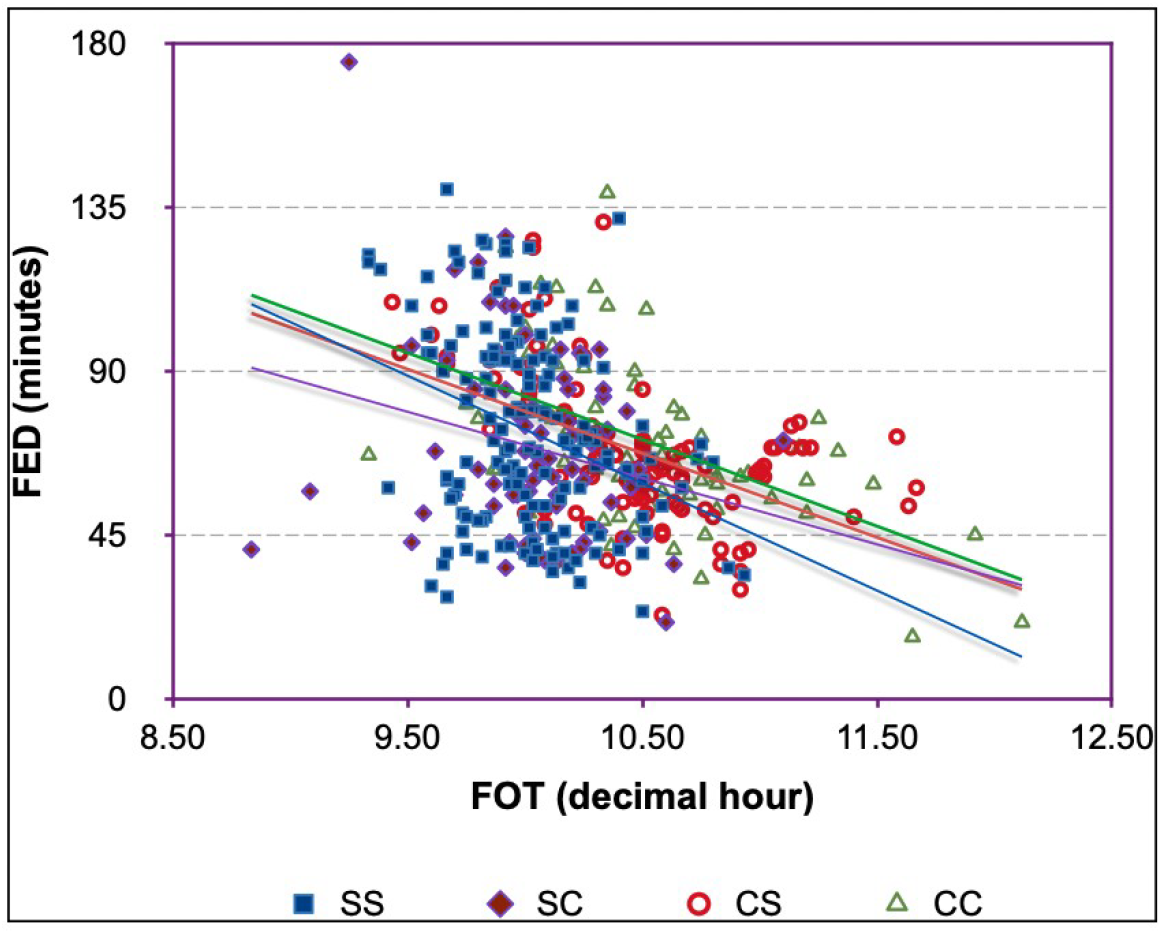
Relationship of FED (minutes) with FOTd at Different Sunlight Exposures during Flower Opening and Closing Time of 420 Landrace Populations (see **Table 2**). The Regression slopes of SS (n =161), SC (n = 78), CS (n = 113) and CC (n = 68) populations are : *b* = –29.5, *R*^2^= 0.107 (*t* = 4.26, *p* <0.00001); *b* = – 18.1, *R*^2^ = 0.058 (*t* = 2.11, *p* <0.05); *b* = – 23.1, *R*^2^ = 0.257 (*t* = 5.76, *p* <0.0001); and *b* = – 23.83, *R*^2^ = 0.27 (*t* = 4.60, *p* <0.00005), respectively.

All the above analyses show the relationship of FED with FOT and SAD of different landraces flowering during short-day season of 2022. The trend of SAD and FED appears to have an inverse association, as corroborated in **Fig. 4**.

### 3.2. Results of the experiment

The records of FOT, FCT, SAD and the sunlight intensity of each of 25 landraces exposed both to direct sunlight and screened from sunlight under shade (mimicking cloudy daylight condition, C) are presented in **Supplementary Table S1**. Our experiment with a set of landraces exposed simultaneously to direct sunlight (denoted hereafter as OL) and under constant shade (denoted as SH) afforded the possibility of finding a definitive relationship of FOT and FED with sunlight intensity. The rice panicles of the ‘population SH’ received only diffused light from two sides, so the range of Lmax under shade (simulating the cloudy condition, C) was at least 2 orders of magnitude lower (min. 100 lux, max. 4568 lux) than the Lmax of sunlight incident on the ‘population OL’ (min. 41.3 K lux, max. 149.2 K lux) of each landrace examined. As the population SH was kept under shade throughout the flowering period, the florets of this group opened as well as closed in the dark, under shade. Thus, the comparison between these two groups essentially compares the effect of SS in the ‘population OL’ on SAD and FED with that of CC condition of the ‘population SH’ under shade.

#### 3.2.1. The effect of open sunlight and shade on SAD and FED

Our experiment with the 25 populations exposed to sunlight (OL) showed a median SAD of 273 min. (IQR = 38) and 302 min. (IQR = 78) under shade (SH, mimicking cloudy condition, C). The distributions of SAD values in the two subpopulations are significantly different (Wilcoxon-Mann-Whitney *z* = 2.62, Bonferroni adjusted *p* < 0.02), as shown in **Fig. 6A**. Conversely, the median FED of the OL group was 71 min (IQR = 16), while that of the SH group was 75 min. (IQR = 31.5). However, as **Fig. 6B** shows, the median FED of the OL group was 70 min (IQR = 13), while that of the SH group was 69 min. (IQR = 21.5). Wilcoxon-Mann-Whitney test cannot detect any difference in the range of FED between the OL and SH groups (*z* = 0.22, Bonferroni adjusted *p* > 0.99).

**Fig 6:**
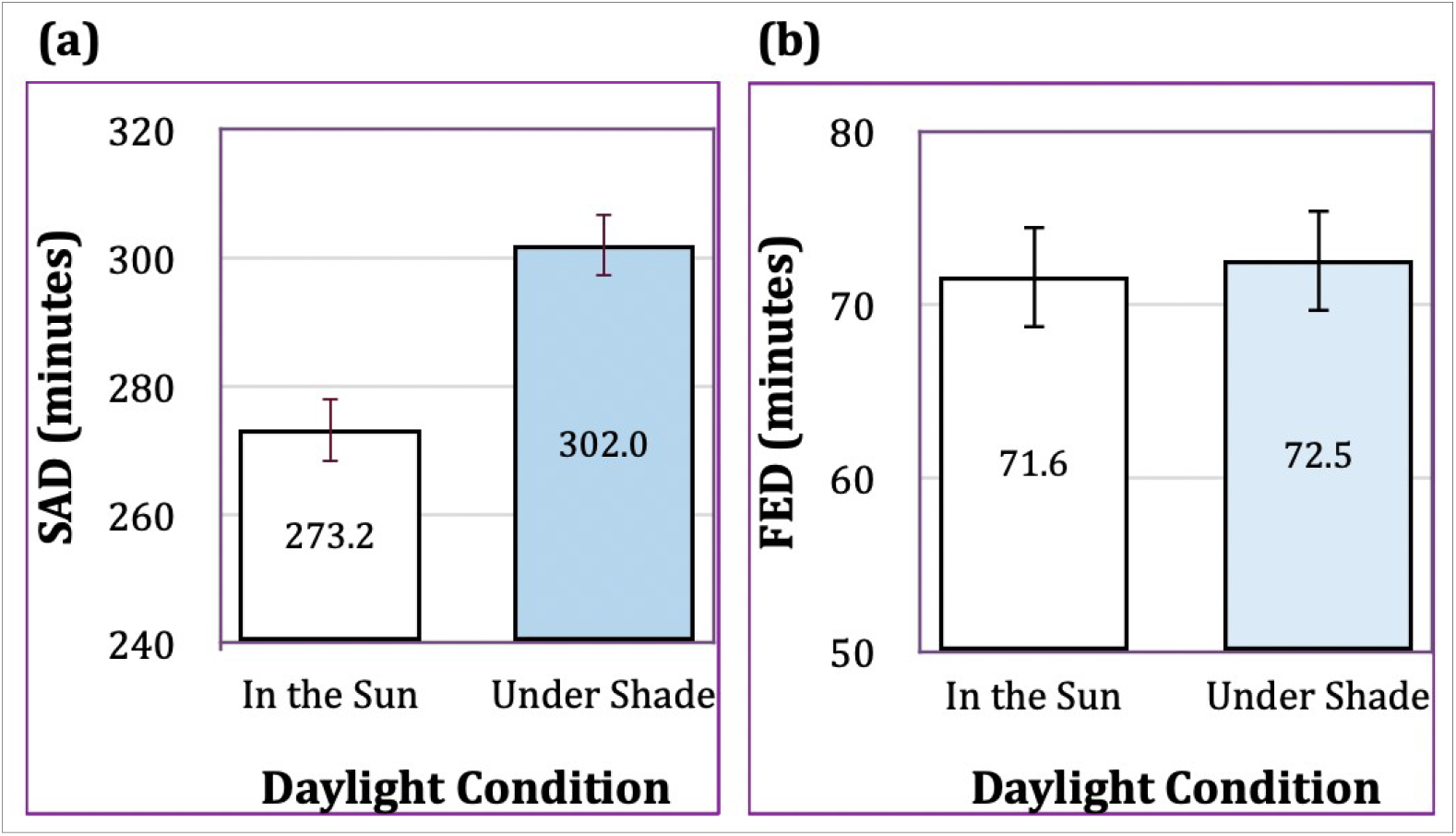
Mean Values of **(a**) SAD and **(b)** FED of Two Populations of 33 Landraces Flowering under Two Different Daylight Conditions. Vertical bars indicate standard errors.

#### 3.2.2 Trends of SAD and FED of landraces opening and closing flowers under the same light conditions

The means of SAD and FED of the S and C populations (**Fig. 2**) were obtained from the landrace populations opening their flowers at either S or C conditions, regardless of S or C conditions at FCT. In contrast, all the flowers of 33 populations in the experimental SH group opened as well as closed in the dark, under shade. Therefore, the effect of darkness on SAD and FED in the 33 landrace populations under SH condition can legitimately be compared only to that of the natural CC group consisting of 68 landraces (**Table 2**); likewise, the effect of OL condition of the experimental 33 populations is comparable only to the SS group consisting of 161 populations (**Table 2**). When the mean of SAD and FED of landrace populations under natural SS and CC conditions are plotted alongside the experimental groups under OL and SH conditions (**Fig. 7a and 7b**), the differences between the groups are evident. To assess the significance of differences between means (*D*) across SS, CC, OL, and SH, we applied ANOVA to both SAD and FED datasets. Following a thorough residual analysis, we checked for heteroscedasticity, and detected no violations of the ANOVA assumptions. Tukey’s HSD test for pairwise comparisons between these groups indicated that:

**Fig 7:**
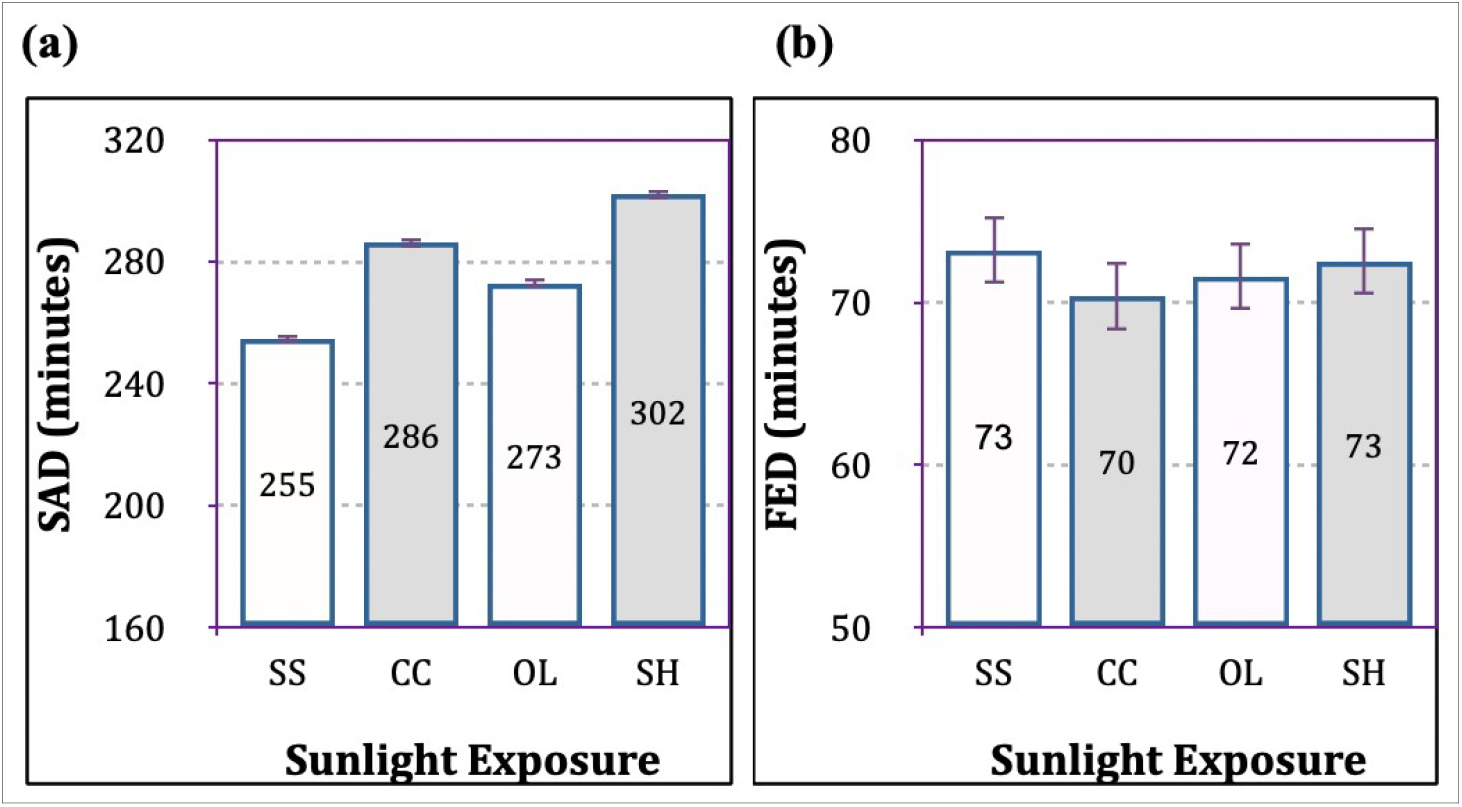
A comparison of **(a)** the mean of SAD and **(b)** the mean of FED of landrace populations that opened and closed flowers during sunny period (SS and OL) with the populations that opened and closed flowers at natural cloudy period (CC) or under shade (SH). The number on the column of each category is the mean value.

a. the mean of SAD of the CC group (*n* = 161) is significantly longer (*D* = 31.47, *p* <0.0001) than that of the SS group (*n* = 68);
b. the mean of SAD of the SH group (*n* = 33) is significantly longer (*D* = 28.79, *p*<0.0001) than that of the OL group (*n* = 33).
c. the difference between means of SAD of the CC group (*n* = 68) under natural cloudy periods and the experimental SH group (*n* = 33) are statistically significant (*D* =−15.79, p *<* 0.03);
d. the means of SAD of the OL group (*n* = 161) is significantly longer (*D* = 18.48, *p* < 0.0001) than that of the SS group (*n* = 33);
e. the difference between means of FED is not significant (*p* >0.9) for any pair of groups.

Our analyses reveal a definitive trend of lengthening of SAD (delayed FOT) under low (<40 K lux) illuminance compared to direct sunlight exposure. This finding is further affirmed by **Fig. 8a**, and a decisive permutation test with 10,000 iterations (**Table 3**). By contrast, FED is confirmed to remain unaffected by the incident sunlight intensity during anthesis (**Fig. 8b** and **Table 3)**.

**Fig 8:**
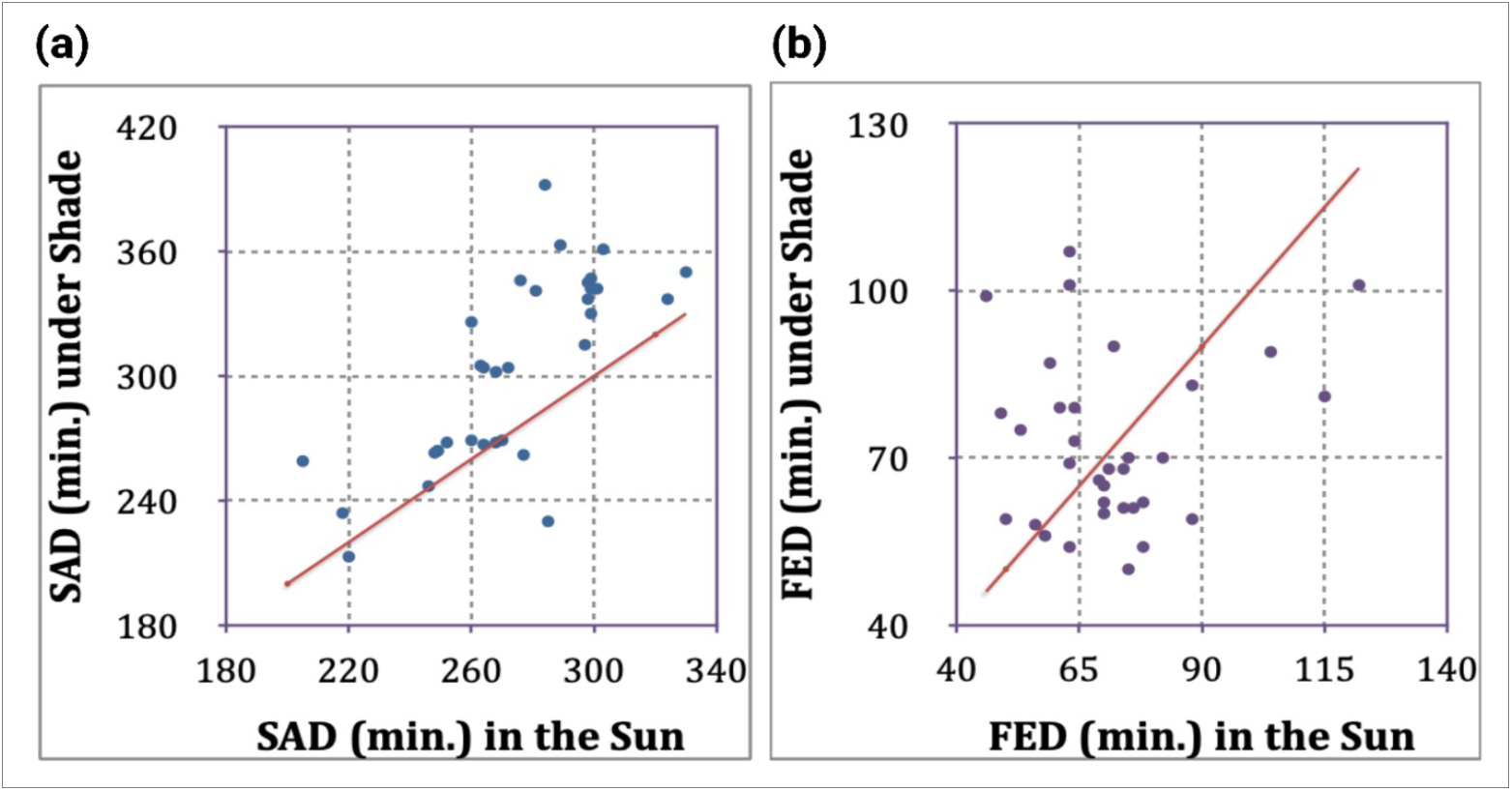
A Comparison of **(a)** SAD and of **(b)** FED of 33 Landrace Populations Flowering in Open Sunlight and Under Shade. The red line in each panel is the isocline.

#### 3.2.3. The relationship between SAD and FED in open sunlight and under cloud cover

The regression of FED on SAD for the S populations shows a strong (*p* <0.0005) inverse relationship (**Fig. 7A**), but the relationship between FED and SAD for the C populations (**Fig. 7B**) is not statistically significant. Furthermore, the extent of change in the FED is not proportionate to the change in SAD. The % difference in SAD between S and C conditions of the same landraces does not show any unimodal correspondence with the respective difference (%) in FED (**Fig. 8**) of the landraces, nor is the relationship significantly different from zero (*R*= 0.03, *t* = 1.17, *p* >0.1).

#### 3.2.4. The relationship between SAD and FED in open sunlight and under experimental shade

The regression of FED on SAD for the OL populations shows a strong (*p* <0.01) inverse relationship (**Fig. 9a**), but the relationship between FED and SAD for the SH populations (**Fig. 9b**) is not statistically significant (*p* >0.4). Furthermore, the extent of change in the FED is not proportionate to the change in SAD. The % difference in SAD between OL and SH subpopulations of the same landraces does not show any unimodal correspondence with the respective difference (%) in FED (**Fig. 10**) of the landraces, as the relationship is almost no different from zero (*R*^2^= 0.007, *t* = 0.47, *p* >0.6).

**Fig 9:**
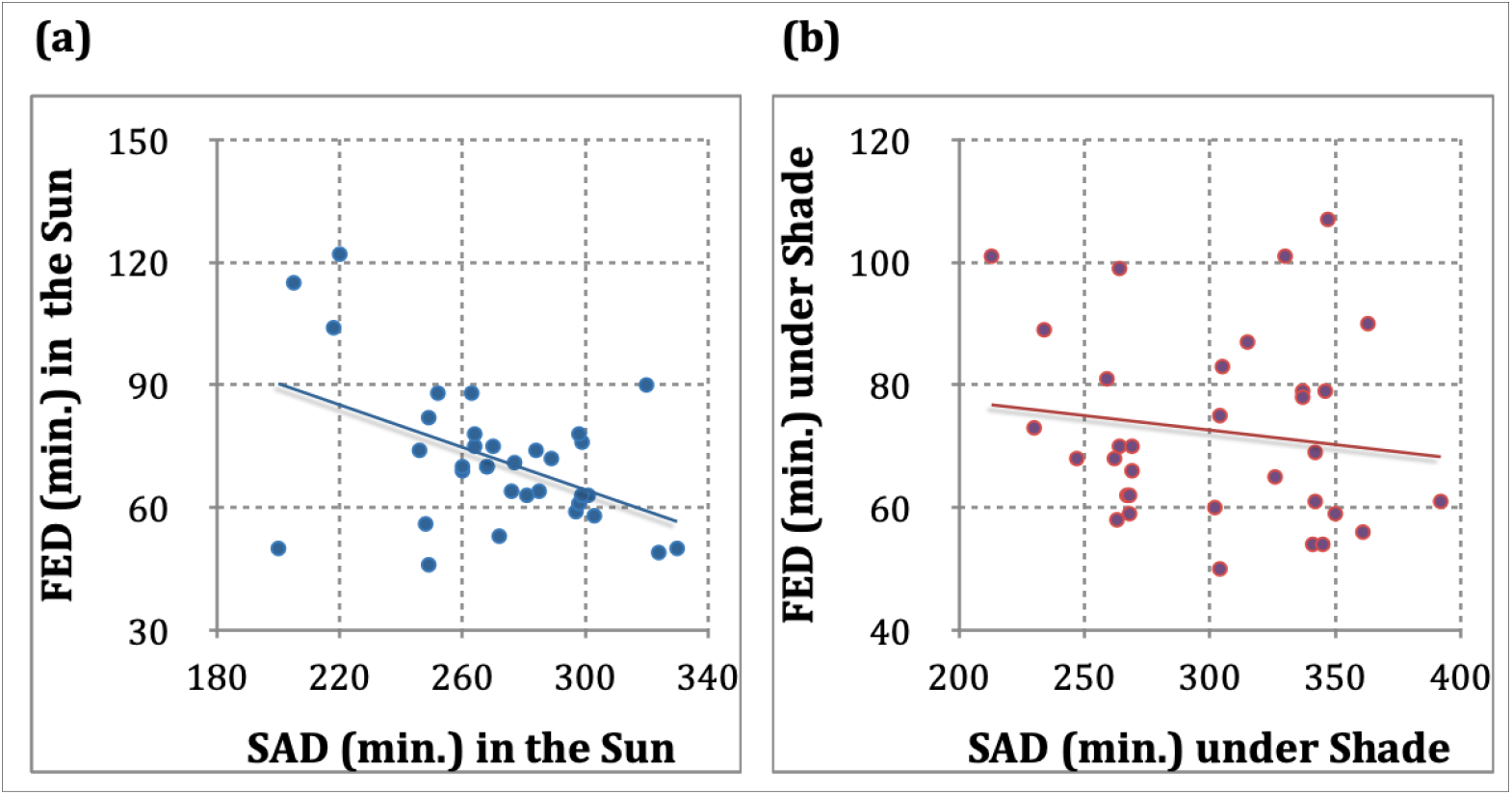
Regression of FED on SAD of of 33 landrace populations **(a)** in open sun (*b* = −0.26, *R*^2^ = 0.22, *t* = 2.82, *p* <0.01) as well as **(b)** under shade (*b* = −0.047, *R*^2^ = 0.02, *t* = 0.80, *p* >0.2.

**Fig 10:**
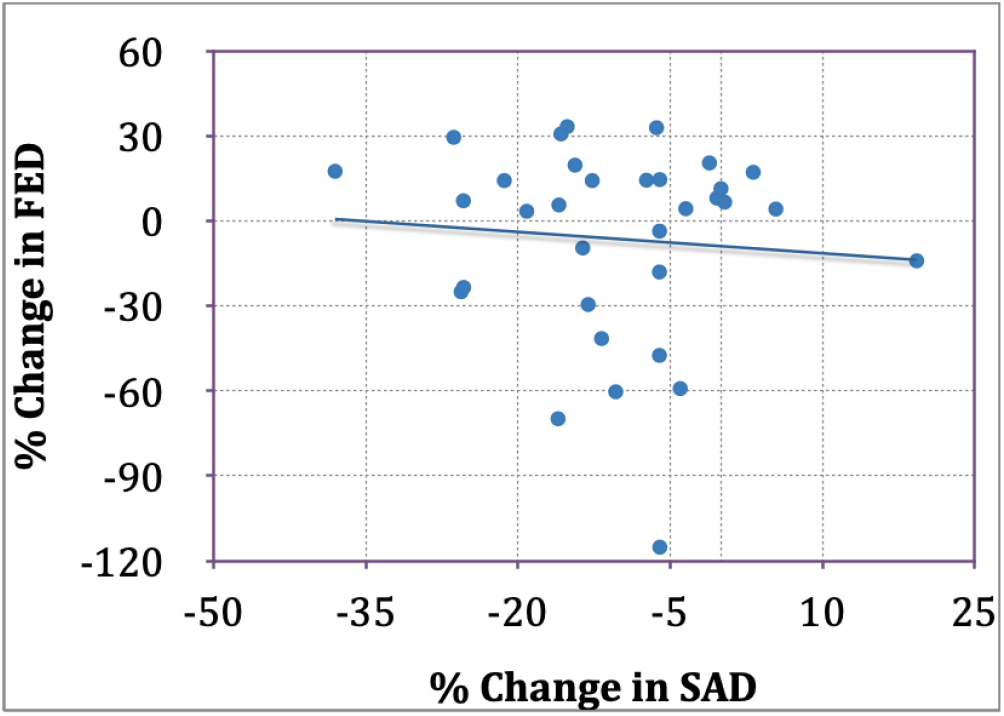
Regression of % change in FED on % change in SAD of 33 populations between OL and SH conditions of daylight at FOT (*b* = – 0.25, *R*^2^ = 0.007, *t* = 0.47, *p* > 0.6).

## 4. Discussion and Conclusions

Some generalizations may be drawn on the above findings, as described below.

**4.1**. In the present study, conducted during the short day (*Aman*) season, rice florets were found to open significantly later during cloudy hours at FOT than when the flowers open in full sun (**Fig. 2A**). This is in agreement with our previous study^[2]^ showing a strong lengthening trend of SAD and shortening duration of FED among 1660 populations. In that previous study we only recorded “cloudy” (C) periods with considerable cloud cover at FOT, although such descriptions included cloudy periods with intermittent sunny intervals. In this study, we have more objectively classified the C period only when the sunlight intensity was recorded to be consistently <40 K lux around the time of opening of the first flowers, and distinguished 181 C populations (receiving <40 K lux at FOT) from 239 S populations (receiving ≥ 40 K lux), comprising a subset of the 1660 populations in our previous study.^[5]^ This reclassification of C and S periods based on the threshold sunlight intensity of 40 K lux, is likely to have obviated any possible misidentification of some populations in C and S categories in the previous study. However, the consonance of the overall pattern of the influence of the S and C periods on both SAD and FED in the previous study with that in the present study implies that there is no Simpson’s paradox, in which the association of variables in a large data set may be reversed in one or more subsets ^[15]^. Excepting 46 repetitions, the populations examined in S and C conditions did not belong to the same landraces. An examination of the same set of landrace populations in both S and C conditions would therefore be more conclusive. Our experimental study of two subpopulations of each of 25 landraces (with 8 repetitions) exposed to full sun and under shade at anthesis corroborates our general observation during S and C periods at anthesis that SAD tends to significantly lengthen on overcast days (when solar illuminance is considerably low (<40 K lux).The FED is inversely related with FOT and SAD on both sunny and cloudy days, and this relationship is strong at *p* < 0.0001 (**Fig. 3**), corroborating our previous finding ^[5]^ using 1114 landraces, including the 388 considered here.
**4.2**. The inverse relationships of FOT and SAD with FED hold unaltered, irrespective of the sunlight intensity at FCT (**Fig. 4**), indicating that the sunlight intensity at the time of closure of florets is not crucial as it is at FOT. However, when the same landraces are examined in open sunlight (OL) and experimentally under shade (SH), the association appears weak (*p* > 0.05) for the C populations (**Fig. 8B**).
**4.3**. The difference of the mean (and median) of SAD between 33 experimental subpopulations of the same 33 landrace populations under open sunlight and under shade is strongly significant, whereas the difference between the medians of FED under OL and SH conditions is not significant (**Table 3**, part **B**).
**4.4.** The % change in FED (min) from open sunlight to considerably reduced illuminance is not proportionate with that of SAD from OL to SH conditions of the same landrace populations, as reflected in a weak regression (p >0.5) of FED on SAD in the experimental populations (**Fig. 9**). This weak association of corresponding changes in SAD and FED under shade appears to confirm that a drastic experimental reduction in illumination on florets of the same landraces does not show any significant effect on FED (**Fig. 5B**). **Fig. 6** further confirms that the influence of low solar illuminance under both natural cloudy hour (C) and artificial shade (SH) on FED is very weak; while the same conditions strongly elicit definitive lengthening of SAD.
**4.5**. The landrace populations exposed to reduced illuminance under natural cloud cover (CC) at both FOT and FCT and under experimental shade (SH) show a similar lengthening of SAD compared to the groups exposed to full sun (SS and OL). **Figures 2a, 6a** and **7a** seem to indicate that SAD of rice landraces is strongly (*p* <0.01) influenced by open sunlight, and is ubiquitously shorter under high (≥ 40 K lux) illuminance than under considerably poor illuminance − both on overcast days and under artificial shade, while the effect of low illuminance on FED is weak (*p* >0.1).
**4.6.** The above findings enable us to present two following explanatory hypotheses.
**4.6.1**. The disappearance of the inverse relationship between SAD and FED under shaded condition (**Fig. 9B**) is a result of the blocking of direct solar illuminance to which rice florets are sensitive. When unshaded, rice florets can receive polarized solar radiation even on cloudy days, which may elicit an appropriate floret closure response, so that a delayed FOT is associated with a shorter FED (**Fig. 3**; also see ref. [5]). In contrast, under experimental shade, the open rice florets are deprived of direct solar cues, resulting in loss of effect of shading on FED.
**4.6.2**. The unequivocal delay in FOT (or, lengthening of SAD) in deficient daylight condition (Fig. 2A, Fig. 3, Fig. 6A and Fig. 7A) is likely an adaptation of the rice plant in anticipation of rainfall, for protection of the pollen from rainwash. This adaptive response of the rice florets to rainfall is further indicated by the facts that (a) rice florets hurry to close immediately after the first raindrop falls on the panicle (pers. observation), and that (b) on a rainy day, FOT is delayed till afternoon [6], even to the next morning (pers. observation). Evidence from a large number of angiosperms [16] confirms that “during the rainy season temporary floral closure would provide protection of the pollen against moisture due to rain, fog, or dew” (p. 5751). After a sufficient delay, when it does not shower, the rice florets begin to open. The lengthening of SAD in experimental SH group, compared to the OL group of populations is similar to the shortening of FED when air temperature is high and approach the maximum day temperature ^[8]^. Just as heat-induced temperature lengthening of SAD and the shortening of FED appears to be an adaptation of the rice plant to protect the pollen from desiccation, the lengthening of SAD at low Lmax on an overcast day (portending rainwash) has an adaptive function to protect the pollen from rainwash. This study provides the first experimental evidence that a delayed FOT in rice is induced by a drastic reduction in incident sunlight intensity on overcast days during short day season, plausibly in order to protect the viable pollen from impending rainwash. We can assert here, paraphrasing van Doorn and Kamdee^[15]^, that “the data do provide suggestions for further experimental work,” which we have presently undertaken.

## List of Abbreviations

FCT: Flower closing time
FD: (First) flowering date
FED: Flower exposure duration
FOT: Flower opening time
IQR: Interquartile range
Lmax: Maximum illuminance (in lux)
SAD: Sunrise-to-anthesis duration

## 5. Supplementary Information

**Fig. S1** (box plot) and **Fig. S2 (**QQ plot) show the near-normal distribution of SAD under sunny and cloudy conditions. A summary of phenological data of 25 landraces populations (with 8 repetitions) experimentally kept under shade and exposed to open sun are given in **Supplementary Table S1**.

## 6. Authors’ contributions

The authors confirm contribution to the paper as follows: study conception and design: Author D.D.; data collection (first and last name): Author D.D., Author M.N.; analysis and interpretation of results: Author SD, Author D.D; draft manuscript preparation: Author D.D. All authors reviewed the results and approved the final version of the manuscript.

## 7. Acknowledgements

We are grateful to Debdulal Bhattacharya for his diligent assistance in recording rice phenological and illuminance data throughout the study period. We are indebted to Dr. Ayan Paul for conducting a part of statistical analyses, and to Mr Avik Saha for his support to our field study. We are grateful to two reviewers and especially to an anonymous Associate Editor for immensely helpful comments and suggestions on an earlier draft of the mss. This study received no institutional funding. We are grateful to Mr Avik Saha for bearing all expenses for field experiments.

## 8. Conflict of Interest

Authors declare that they have no conflict of interest.

## 9. Data availability

The primary data pertaining to FOT, FED and SAD of 1660 landrace populations cultivated from 2020 to 2022 are openly available from Harvard Dataverse^[9]^. The records of illuminance during anthesis of 390 rice landraces examined here are available from Harvard Dataverse^[14]^. A summary of the data pertaining to our experiment with 25 landrace populations are given in **Supplementary Table S1**.

## Notes

### Competing Interest Statement

The authors have declared no competing interest.

### Summary of Updates

The Lmax incident on the population OL during the study period always exceeded 30000 lux. Therefore, the threshold illuminance in cloudy hours was set at 30 K lux (instead of 20K lux used earlier).

