## Supplementary Table S1 for "The Effect of Sunlight Intensity on Flower Opening Time and Exposure Duration in Rice (*Oryza sativa* ssp. *indica*) Landraces"

**Table S1**: Flower Opening Time (FOT), Flower Exposure Duration (FED), Sunrise-to-Anthesis Duration (SAD) of 25 Landraces (33 Populations), and Maximum Illuminance (lux) measured at FOT of Panicles Exposed to Full Sunlight and under Shade.

| **Land race** | **Date of Anthesis** | **In Open Sun (OL)** | | | | **Under Shade (SH)** | | | |
| --- | --- | --- | --- | --- | --- | --- | --- | --- | --- |
|  |  | **FOT** | **SAD**  **(min.)** | **FED**  **(min)** | **LMAX (lux)** | **FOT** | **SAD**  **(min.)** | **FED**  **(min)** | **LMAX (lux)** |
| BAID DULAH | 06/10/22 | 10:32 | 285 | 64 | 104260 | 09:37 | 230 | 73 | 100 |
| BANKULI | 30/10/22 | 10:58 | 303 | 58 | 59770 | 11:56 | 361 | 56 | 1122 |
| BANKULI | 30/10/22 | 10:52 | 297 | 59 | 58900 | 11:10 | 315 | 87 | 1469 |
| BHASA KALMI | 09/11/22 | 11:24 | 324 | 49 | 124100 | 11:37 | 337 | 78 | 309 |
| BHONDA DHAN | 06/11/22 | 10:57 | 298 | 78 | 116990 | 11:44 | 345 | 54 | 352 |
| DUDHE BOLTA | 15/10/22 | 10:14 | 264 | 78 | 58830 | 10:17 | 267 | 62 | 195 |
| DUDHE BOLTA | 15/10/22 | 10:02 | 252 | 88 | 60400 | 10:18 | 268 | 59 | 673 |
| GITA | 02/11/22 | 10:38 | 281 | 63 | 95160 | 11:38 | 341 | 54 | 299 |
| GITA | 02/11/22 | 10:25 | 268 | 70 | 144900 | 10:59 | 302 | 60 | 789 |
| HALDÉ BATALI | 01/11/22 | 10:40 | 284 | 74 | 95690 | 12:28 | 392 | 61 | 244 |
| HALDÉ BATALI | 01/11/22 | 10:45 | 289 | 72 | 72680 | 11:59 | 363 | 90 | 408 |
| HINCHÉ SAROO | 06/11/22 | 10:57 | 298 | 61 | 70250 | 11:36 | 337 | 79 | 239 |
| KALABATI | 27/10/22 | 10:30 | 276 | 64 | 96910 | 11:40 | 346 | 79 | 2458 |
| KAMATH | 06/11/22 | 10:58 | 299 | 76 | 62270 | 11:41 | 342 | 61 | 403 |
| KARPURTUL | 19/10/22 | 10:15 | 264 | 75 | 112020 | 10:55 | 304 | 50 | 463 |
| KBA LYNGKOT | 02/11/22 | 10:20 | 263 | 88 | 125250 | 11:02 | 305 | 83 | 320 |
| KETAKI | 29/10/22 | 10:54 | 299 | 63 | 102000 | 11:25 | 330 | 101 | 958 |
| KONG HING | 02/10/22 | 09:56 | 249 | 46 | 129390 | 10:11 | 264 | 99 | 635 |
| MISE BATTA | 11/11/22 | 11:00 | 299 | 63 | 83880 | 11:48 | 347 | 107 | 301 |
| NIROJA | 06/11/22 | 11:00 | 301 | 63 | 73910 | 11:41 | 342 | 69 | 768 |
| PAYJAM | 16/10/22 | 09:59 | 249 | 82 | 127600 | 10:14 | 264 | 70 | 228 |
| PAYJAM | 16/10/22 | 10:10 | 260 | 69 | 103620 | 10:19 | 269 | 66 | 174 |
| PATNAI | 17/10/22 | 10:19 | 268 | 70 | 52940 | 10:19 | 268 | 62 | 248 |
| RELDO | 15/10/22 | 09:28 | 218 | 104 | 77180 | 09:44 | 234 | 89 | 223 |
| RELDO | 14/10/22 | 09:30 | 220 | 122 | 136220 | 09:23 | 213 | 101 | 129 |
| SALEM SANNA | 27/11/22 | 10:30 | 260 | 70 | 105200 | 11:36 | 326 | 65 | 210 |
| SOPUR DHAN | 15/10/22 | 10:20 | 270 | 75 | 50030 | 10:19 | 269 | 70 | 160 |
| SOPUR DHAN | 15/10/22 | 10:27 | 277 | 71 | 103600 | 10:12 | 262 | 68 | 198 |
| SHIYAL RAJ | 07/10/22 | 09:54 | 246 | 74 | 101900 | 09:55 | 247 | 68 | 4568 |
| SHIYAL RAJ | 06/10/22 | 09:55 | 248 | 56 | 111800 | 10:10 | 263 | 58 | 1518 |
| TUPRU | 28/10/22 | 10:26 | 272 | 53 | 73370 | 10:58 | 304 | 75 | 2825 |
| TARMANGA | 27/10/22 | 11:24 | 330 | 50 | 87680 | 11:44 | 350 | 59 | 960 |
| UMURIA CHURI | 16/10/22 | 09:15 | 205 | 115 | 53780 | 10:09 | 259 | 81 | 342 |
