## Supplementary Fig. S1 and S2 for "The Effect of Sunlight Intensity on Flower Opening Time and Exposure Duration in Rice (*Oryza sativa* ssp. *indica*) Landraces"

**Supplementary Figures**

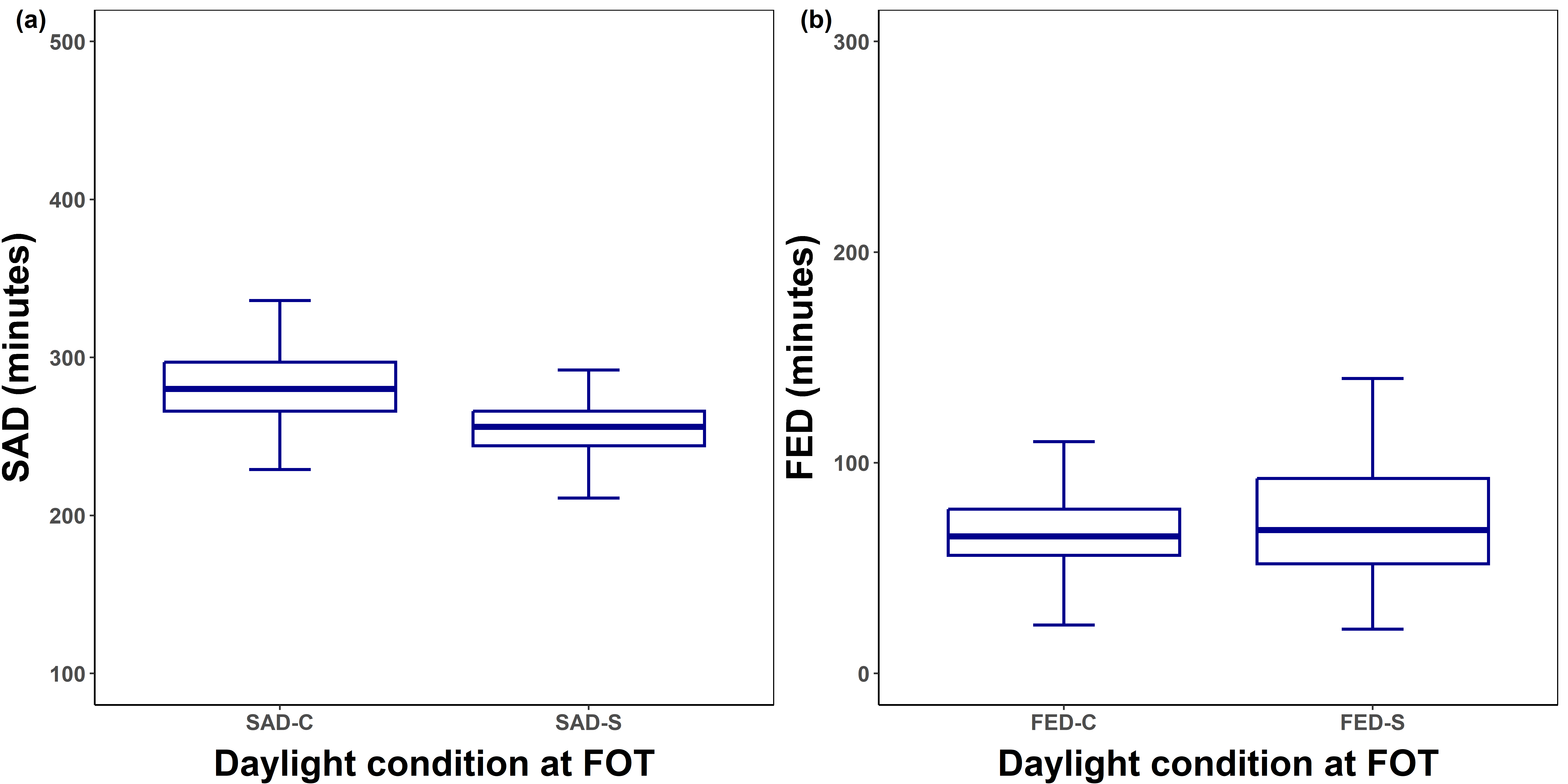

**Fig. S1**: Box Plots of (a) SAD at FOT and (b) FED during Sunny (S) and during Cloudy Period (C).

'

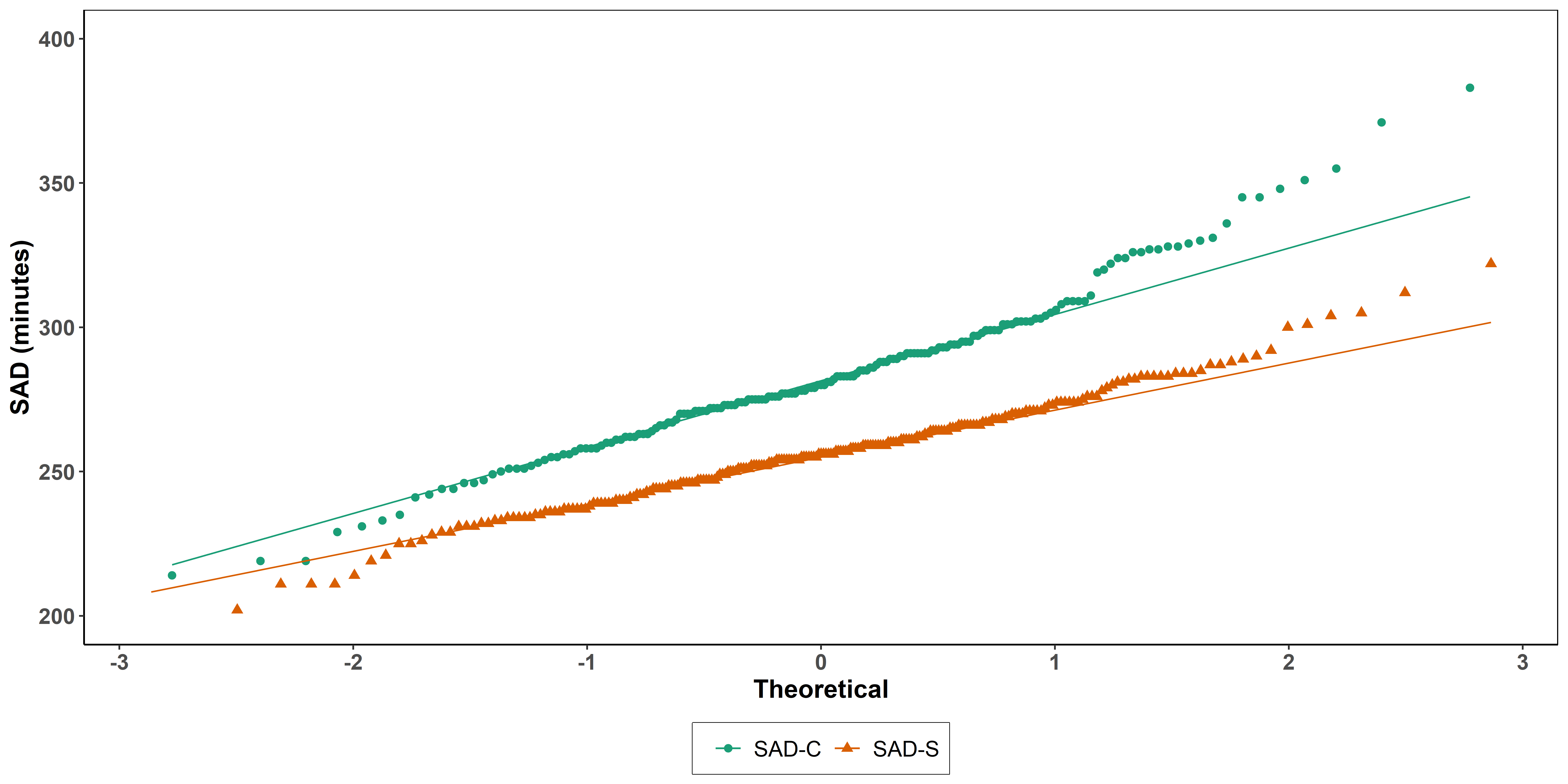

**Fig. S2**: QQ Plots of SAD at FOT during Sunny (S) and during Cloudy Period (C).
